# Deriving C_4_ photosynthesis parameters by fitting intensive A/C_i_ curves

**DOI:** 10.1101/153072

**Authors:** Haoran Zhou, Erol Akçay, Brent R. Helliker

## Abstract

Measurements of photosynthetic assimilation rate as a function of intercellular CO_2_ (*A*/*C*_i_ curves) are widely used to estimate photosynthetic parameters for C_3_ species, yet few parameters have been reported for C_4_ plants, because of a lack of estimation methods. Here, we extend the framework of widely-used estimation methods for C_3_ plants to build estimation tools by exclusively fitting intensive *A*/*C*_i_ curves (6-8 more sampling points) for C_4_ using three versions of photosynthesis models with different assumptions about carbonic anhydrase processes and ATP distribution. We use simulation-analysis, out-of-sample tests, existing in vitro measurements and chlorophyll-fluorescence-measurements to validate the new estimation methods. Of the five/six photosynthetic parameters obtained, sensitivity analyses show that maximal-Rubisco-carboxylation-rate, electron-transport-rate, maximal-PEP-carboxylation-rate and carbonic-anhydrase were robust to variation in the input parameters, while day-respiration and mesophyll-conductance varied. Our method provides a way to estimate carbonic anhydrase activity, a new parameter, from *A*/*C*_i_ curves, yet also shows that models that do not explicitly consider carbonic anhydrase yield approximate results. The two photosynthesis models, differing in whether ATP could freely transport between RuBP and PEP regeneration processes yielded consistent results under high light, but they may diverge under low light intensities. Modeling results show selection for Rubisco of low specificity and high catalytic rate, low leakage of bundle sheath and high PEPC affinity, which may further increase C_4_ efficiency.

## 1. INTRODUCTION

Key photosynthetic parameters allow for the assessment of how biochemical and biophysical components of photosynthesis affect net carbon assimilation in response to environmental changes, phenotypic/genotypic differences, and genetic modification. The changes in net assimilation (*A*_n_) that occur along with the changes of intercellular CO_2_ concentration (*C*_i_) —or *A*/*C*_i_ curves— are widely used to estimate photosynthetic parameters for C_3_ species. In particular, the method by Sharkey et al. (2007), based on the C_3_ photosynthesis model of Farquhar et al. (1980; FvCB model), has been one of the most widely used tools since it is based exclusively on *A*/*C*_i_ curves, which are easy to measure in both lab and field conditions.

Fewer estimates of photosynthetic parameters have been reported for C_4_ species, as there has been a lack of accessible C_4_ estimation methods. Several recent studies, however, used *A*/*C*_i_ curves to estimate photosynthesis parameters based on the C_4_ photosynthesis model of von Caemmerer (2000)(Ubierna et al., 2013; Bellasio et al., 2015). These studies use partial *A*/*C*_i_ curves; measuring assimilation rates for only a few CO_2_ concentrations coupled with ancillary measurements of chlorophyll fluorescence and/or 2% O_2_. While these estimation methods lead to estimates of photosynthetic parameters, the additional measurements they require make estimation more cumbersome for field work or large-scale sampling. Theoretically, it is possible to estimate photosynthetic parameters by exclusively fitting *A*/*C*_i_ curves to a C_4_ photosynthesis model. In this paper, we propose the method to estimate C_4_ photosynthesis parameters using only A/C_i_ curves.

There are several potential problems with *A*/*C*_i_ –based estimation methods for C_3_ plants that carry over to existing C_4_ methods (Gu et al. 2010); it is therefore important to develop a C_4_ estimation method with improvements to solve the general problems and drawbacks outlined below. First, the structure of the FvCB model makes it easy to be over-parameterized. Second, a general shortcoming for the estimation methods is that they require an artificial assignment of the RuBP regeneration and Rubisco carboxylation limitation states to parts of the *A*/*C*_i_ curves (Xu and Baldocchi, 2003; Ethier et al., 2006; Ubierna et al., 2013; Bellasio et al., 2015), which has turned out to be problematic (Type I methods) (Gu et al. 2010). These methods assume constant transition points of limitation states for different species. Furthermore, Type I methods tend to minimize separate cost functions of different limitation states instead of minimizing a joint cost function. Some recent estimation methods for C_3_ species ameliorate these problems by allowing the limitation states to vary at each iterative step of minimizing the cost function (Type II methods;Dubois et al., 2007; Miao et al., 2009; Yin et al., 2009; Gu et al., 2010). However, for these type II methods, additional degrees of freedom in these “auto-identifying” strategies can lead to over-parameterization if limitation states are allowed to change freely for all data points. Gu et al. (2010) also pointed out that existing Type I and Type II methods fail to check for inadmissible fits, which happen when estimated parameters lead to an inconsistent identification of limitation states from the formerly assigned limitation states. More specifically to C_4_, the recently developed C_4_ estimation methods artificially assign limitation states for A/C_i_ curves (Ubierna et al., 2013; Bellasio et al., 2015) and also did not check for inadmissible fits.

We developed methods to estimate photosynthetic parameters for C_4_ species based solely on fitting intensive *A*/*C*_i_ curves to a C_4_ photosynthesis model (von Caemmerer, 2000). The intensive *A*/*C*_i_ curves (*A*/*C*_i_ curves with 6-8 more sampling points than the common *A*/*C*_i_ for C_3_ species) are important for two reasons: First, at low *C*_i_, the slope of *A*/*C*_i_ is very steep and the assimilation rate saturates quickly. Second, C_4_ species have more photosynthetic parameters as the carbon concentrating mechanism adds complexity. Additionally, carbonic anhydrase catalyzes the first reaction step for C_4_ photosynthesis (Jenkins et al., 1989), and it has been commonly assumed to not limit CO_2_ uptake in estimation methods and C_4_ models (von Caemmerer, 2000; Yin et al., 2011b). Recent studies, however, showed evidence of potential limitation by carbonic anhydrase (von Caemmerer et al., 2004; Studer et al., 2014; Boyd et al., 2015; Ubierna et al., 2017).

Therefore, first, we built estimation methods using two different fitting procedures of Sharkey et al. (2007) and Yin et al. (2011b) without considering carbonic anhydrase activity. Then, we add carbonic anhydrase limitation into the estimation method. We can also use this approach to examine how the carbonic-anhydrase-limitation assumption impacts parameter estimation, and whether the modeling of C_4_ photosynthesis can be simplified by omitting it. All together, our method estimates five to six photosynthesis parameters: (1) maximum carboxylation rate allowed by ribulose 1,5-bisphosphate carboxylase/oxygenase (Rubisco) (*V*_cmax_), (2) rate of photosynthetic electron transport (*J*), (3) day respiration (R_d_), (4) maximal PEP carboxylation rate (*V*_pmax_), (5) mesophyll conductance (*g*_m_), and optionally (6) the rate constant for carbonic anhydrase hydration activity (*k*_CA_). These approaches yield the following improvements to eliminate common problems occurring in the previous C_3_ and C_4_ estimation methods: avoiding over-parameterization, maximizing joint cost function, freely determining transition points instead of assigning in advance, and checking for inadmissible fits. Second, since both RuBP regeneration and PEP regeneration need ATP (Hatch, 1987), we also examine two different assumptions about ATP distribution between RuBP regeneration and PEP regeneration in C_4_ photosynthesis models. Third, we validate the estimation methods in four independent ways, using: (i) simulation tests using *A*/*C*_i_ curves generated using our model with known parameters and adding random errors, (ii) out of sample test, (iii) existing in vitro measurements and (iv) Chlorophyll fluorescence measurement. Finally, we used the C_4_ photosynthesis model to perform sensitivity analyses and simulation analyses for important physiological input parameters. These analyses allow us to illustrate the underlying physiological significance of these parameters to the ecology and evolution of the C_4_ photosynthesis pathway.

## 2. MATERIALS and METHODS

### 2.1 C_4_ Mechanism

The CO_2_ concentrating mechanism of C_4_ pathway increases CO_2_ in the bundle sheath cells to eliminate photorespiration. Like the C_3_ pathway, the diffusion of CO_2_ starts from the ambient atmosphere through stomata into intercellular spaces, and then into the mesophyll cells. In the mesophyll cells, the first step is the hydration of CO_2_ into HCO_3_ ^-^ by carbonic anhydrase. PEPC, then, catalyze HCO_3_ ^-^ and PEP into C_4_ acids and the C_4_ acids are transported to the bundle sheath cells. In the bundle sheath cell, C_4_ acids are decarboxylated to create a high CO_2_ environment for the C_3_ photosynthetic cycle, and PEP is regenerated. All the modeling equations and mechanistic processes used for our estimation method are from von Caemmerer (2000), Hatch and Burnell (1990), Boyd et al. (2015) and Ubierna et al. (2017)(Supplementary Methods).

Given the two limitation states of C_4_ cycle (PEP carboxylation (*V*_pc_) and PEP Regeneration (*V*_pr_)), and two limitation states of C_3_ cycle (RuBP carboxylation (*A*_c_) and RuBP Regeneration (*A*_j_)) in the C_4_ photosynthesis model, there are four combinations of limitation states (as Yin et al., 2011b, Fig. 1): RuBP carboxylation and PEP carboxylation limited assimilation (AEE), RuBP carboxylation and PEP regeneration limited assimilation (ATE), RuBP regeneration and PEP carboxylation limited assimilation (AET) and RuBP regeneration and PEP regeneration limited assimilation (ATT). Since the C_4_ cycle operates before the C_3_ cycle and provides substrates for the C_3_ cycle, the determination process of *A*_n_ is as follows:

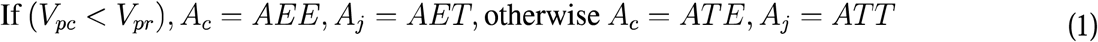

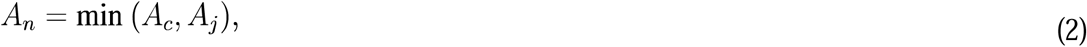

which we used for our estimation method.

**Fig. 1.**
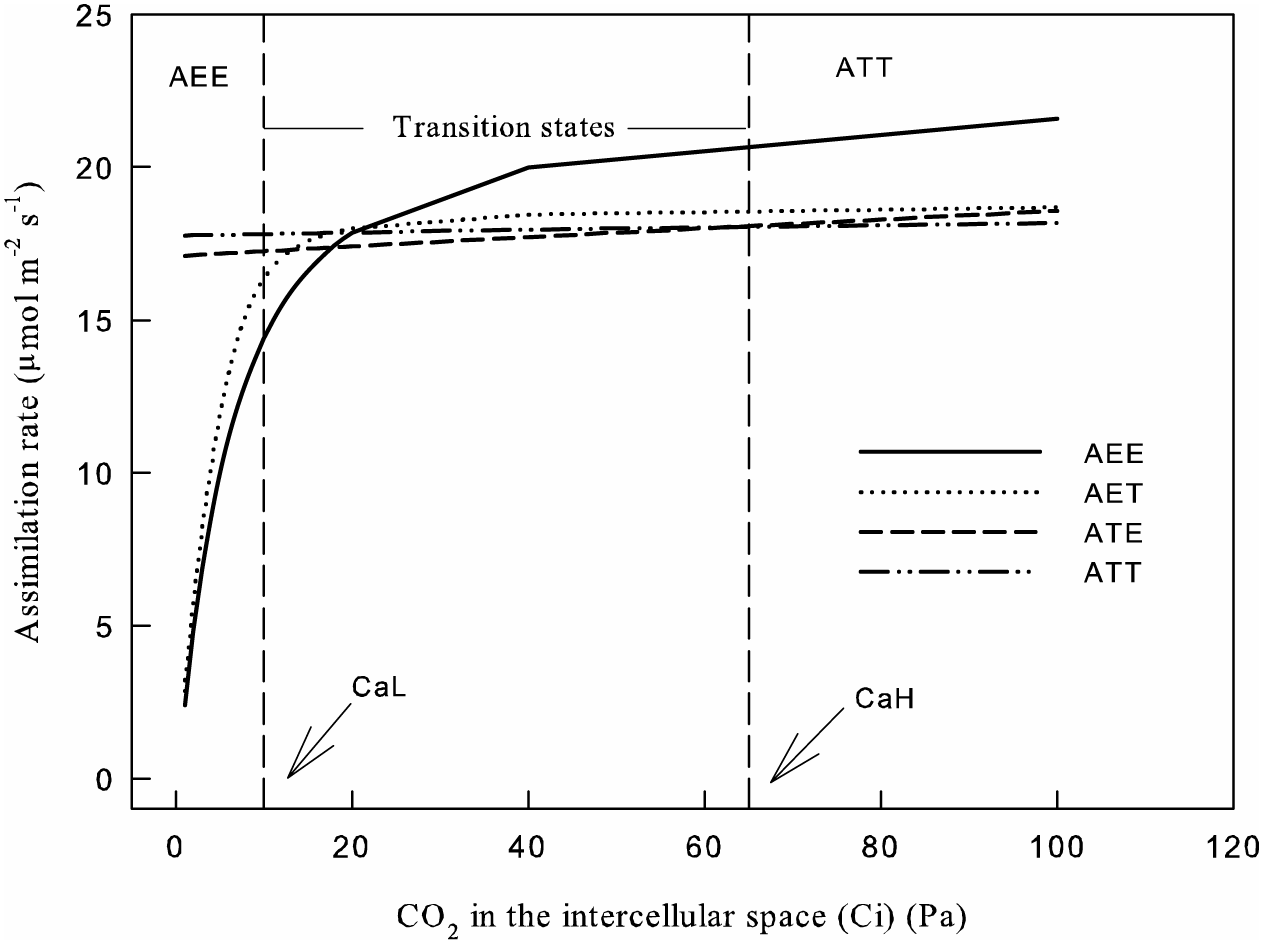
An introduction of how our estimation methods assign transition points between limitation states. AEE represents RuBP carboxylation, and PEP carboxylation limited assimilation rate, ATT represents RuBP regeneration and PEP regeneration limited assimilation rate. Transition states indicate assimilation could be limited by AEE, ATT, ATE (RuBP carboxylation and PEP regeneration) and AET (RuBP regeneration and PEP carboxylation). Our algorithm allows the transition states to be freely limited by the above four conditions from a lower bound (*CaL*, 10 Pa for instance) and a higher bound (*CaH*, 65 Pa for example), indicated by the dashed vertical lines in the figure.

### 2.2 Plant Material

We performed intensive *A*/*C*_i_ curves on nine different C_4_ species to develop and examine the efficacy of our estimation tools: *Zea mays* L., *Eragrostis trichodes* (Nutt.) Alph. Wood, *Andropogon virginicus* L., *Schizachyrium scoparium* (Michx.) Nash, *Panicum virgatum* L., *Panicum amarum* Elliott, *Setaria faberi* Herrm., *Sorghastrum nutans* (L.) Nash and *Tripsacum dactyloides* (L.) L. The intensive *A*/*C*_i_ curves contain more sample points under more CO_2_ concentrations than the default curve used for C_3_ species. Here we set the CO_2_ concentrations as 400, 200, 50, 75, 100, 125, 150, 175, 200, 225, 250, 275, 300, 325, 350, 400, 500, 600, 700, 800, 1000, 1200, 1400 ppm under light intensity of 1500 μmolm^-2^s^-1^ (light intensity encountered by the plants in greenhouse). At each point, data was recorded when the intercellular CO_2_ concentration equilibrated within 2-5 minutes. The datasets were obtained using a standard 2 × 3 cm^2^ leaf chamber with a red/blue LED light source of LI-6400 (LI-COR Inc., Lincoln, NE, USA). If the stomatal conductance of a species does not decrease quickly at high CO_2_, then the sample points at the high CO_2_ level can be increased. Fluorescence was measured along with *A*/*C*_i_ curves for seven C_4_ species (CO_2_ concentration is similar with above). After each change of CO_2_ concentration and *A* reached steady state, the quantum yield was measured by multiphase flash using a 2 cm^2^ fluorescence chamber head (Bellasio et al., 2014). All the measurements are conducted at 25°C and VPD is controlled at 1-1.7kPa. The cuvette was covered by Fun-Tak to avoid and correct for the leakiness (Chi et al., 2013).

### 2.3 Estimation Protocol

We implemented the estimation methods using the non-linear curve-fitting routine in MS Excel (Supplementary Material I, II, III) and independently in R (“C4Estimation”) to get solutions that minimize the squared difference between observed and predicted assimilation rates (*A*). Five (or six when considering carbonic anhydrase) parameters will be estimated by fitting the *A*/*C*_i_ curve: *V*_cmax_, *J*, *R*_d_, *V*_pmax_, *g*_m_, and *k*_CA._ Other input parameters for C_4_ are in Table S1.

#### Input data sets and preliminary calculations

The input data sets are the leaf temperature during measurements, atmosphere pressure, two CO_2_ bounds (*CaL* and *CaH* discussed in the following section), and the assimilation rates (*A*) and the *C*_i_s (in ppm) in the *A*/*C*_i_ curve. Also, reasonable initial values of output parameters need to be given in the output section to initiate the non-linear curve fitting (Supplementary Material IV). *C*_i_ will be adjusted from the unit of ppm to the unit of Pa inside the program as suggested by Sharkey et al. (2007).

#### Estimating limitation states

We set upper and lower limits to the value of *C*_i_ between which the assimilation rates are freely determined by limitation states. Also, we can avoid over-parameterization by pre-assigning limitation states at the lower and upper ends of the *C*_i_ range. We assumed that under very low *C*_i_ (*CaL*), CO_2_ is the limiting substrate; thus, *V*_p_ is limited by *V*_pc_ and *A* is given by *A*_c_ (AEE); under very high *C*_i_ (*CaH*) electron transport is limiting, thus, *V*_p_ is limited by *V*_pr_ and *A* is given by *A*_j_ (ATT) (Fig. 1). The points between *CaL* to *CaH* are freely determined by AEE, ATE, AET or ATT from eq. (16) and (17) to minimize the cost function. We suggest setting *CaL* as 10 Pa initially, then adjusting based on the preliminary results. The points of constant *A* at high *C*_i_ end can initially be set as being limited by ATT primarily (based on the three points, we can *CaH*) or use 65 Pa as the first trial. The range of freely determined points can be adjusted by users by setting appropriate *CaL* and *CaH*. In the column of “Estimate Limitation”, whether the data points are limited by AEE (represented by “1”), ATT (represented by “4”) or freely vary (represented by “0”), all the assignments of “1”, “4” and “0” are determined automatically by the given values of *CaL* and *CaH*. One can input “-1” to disregard a data pair. Users can adjust limitation states according to how many points and the range of *C*_i_ they have in their *A*/*C*_i_ curves.

We assume different processes in the C_4_ photosynthesis are coordinated with each other and co-limit the assimilation rate (Sharkey et al., 2007; Yin et al., 2011b; Ubierna et al., 2013; Bellasio et al., 2015). Thus, the estimation parameters allow the limitation states to be compactly clustered with each other (Fig. 1). However, if there were only a few points under *CaL*, the estimation results will depend heavily on the given initial values and unbalanced results would be more likely. Fig. S1 shows an example of unbalanced estimation results by deleting some points under 10 Pa or setting a very low *CaL*: in the estimation results, *A*_n_ is limited by *AEE* at very low *C*_i_ and is mostly limited by *A*_j_ (shown by AET and ATT) in the C_3_ cycle. In this case, *A*_c_ (shown by AEE and ATE) has a clear redundancy at higher *C*_i_. Unbalanced results happened when there are not enough constraints points under *CaL* or above *CaH*. Such results explain why intensive *A*/*C*_i_ curves are preferred, especially more measuring points under the lower end and higher end of *C*_i_. However, existing *A*/*C*_i_ data with 14 points might be used in the current estimation method if there are at least four points below *CaL* and three points above *CaH*.

#### Estimation algorithm and fitting procedures

The objective of our estimation methods is to minimize the following joint cost function (eq. 3 and 4) by varying the above five or six output parameters (*V*_cmax_, *J*, *R*_d_, *V*_pmax_, *g*_m_, and *k*_CA_):

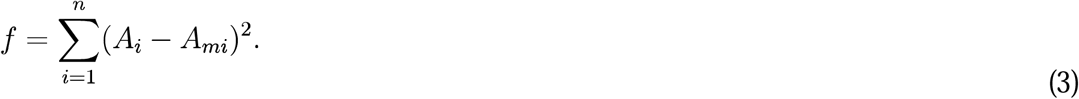

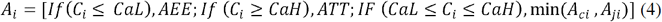

*n* is the total number of observations, *A*_ci_ is determined by AEE and ATE and *A*_ji_ is determined by AET and ATT from eq. (1), *A*_mi_ is the observed net assimilation rate. In this calculation, we take Michaelis-Menten constant of Rubisco activity for CO_2_ (*K*_c_), Michaelis-Menten constant of Rubisco activity for O_2_ (*K*_o_), the specificity of Rubisco (*γ*\*), Michaelis-Menten constants of PEP carboxylation for CO_2_ or HCO3^-^ (*K*_p_), the fraction of O_2_ evolution occurring in the bundle sheath (*α*) and bundle sheath conductance (*g*_bs_) as given (input parameters), similar to Sharkey et al. (2007). We conduct further sensitivity analyses in the following section to determine the effects of variability of these inputs parameters on the estimation results.

We used two fitting procedures in the current study: one was from Sharkey et al. (2007), which is an implicit minimization of error (Supplementary Material I, III), and the other one was based on the explicit calculations given by Yin et al. (2011b)(Supplementary Material II). For the method of Sharkey et al. (2007), “estimated” *A*_n_ was calculated using the above equations and observed *A*_n_ values. We call them “estimated”, because when we calculate *A*_n_, observed *A*_n_ is used to calculate intermediate parameters, for example, the CO_2_ concentration in mesophyll cells (*C*_m_), the CO_2_ concentration in bundle sheath (*C*_bs_), which we then use to calculate *A*_c_ and *A*_j_. The objective function is to minimize the sum of square errors between “estimated” *A*_n_ and observed *A*_n_ (Simulation Error in Supplementary Material I, III). For the model without carbonic anhydrase, Yin et al. (2011b) gave explicit solutions for AEE, ATE, AEE, and ATT). “Explicit” here means the assimilation rates are totally calculated by the estimated parameters without calculating the intermediates with observed A_n_. These calculations give us the real estimation error of our fitting procedure for models without carbonic anhydrase and thus provide a validation for the goodness of fit (“True Error” in Supplementary Material I-III).

#### Checking inadmissible fits

We made it possible to check the inadmissible fits for limitation states in our estimation method. After the estimation process finishes, the limitation states based on the estimated parameters will be calculated in the last column. If the calculated limitation states are inconsistent with the assigned ones in the estimation method, one needs to readjust the assignment of the “Estimate Limitation” (adjust *CaL* or *CaH*) and rerun the estimation method, until they are consistent with each other.

## 3. RESULTS

### 3.1 Estimation results and assumptions

Estimation methods based on assumptions with and without carbonic anhydrase yield similar results (Supplementary material V). In Supplementary material III, carbonic anhydrase indeed shows limitation to *V*_pc_, which confirms its potential role as a limiting step in the C_4_ cycle. However, *V*_pc_ calculated from CO_2_ are only a little higher than *V*_pc_ calculated from HCO_3_^-^, which resulted in the similar estimation results. In addition, the estimation errors and true errors from Yin’s equations are quite small (average<1), and also similar between models with and without carbonic anhydrase.

Estimation methods based on the two equations of different assumptions about electron transport between RuBP regeneration and PEP regeneration yield consistent parameter estimates and assimilation- CO_2_ response curves (Fig. 2), but there were minor differences. The second assumption that ATP, resulting from electron transport, is freely allocated between PEP carboxylation-regeneration and RuBP regeneration leads to a bump at low CO_2_ when estimating ATE. The two assumptions produce different ATE under low CO_2_; but this is largely inconsequential because, under low CO_2_, assimilation is usually limited by AEE.

**Fig. 2.**
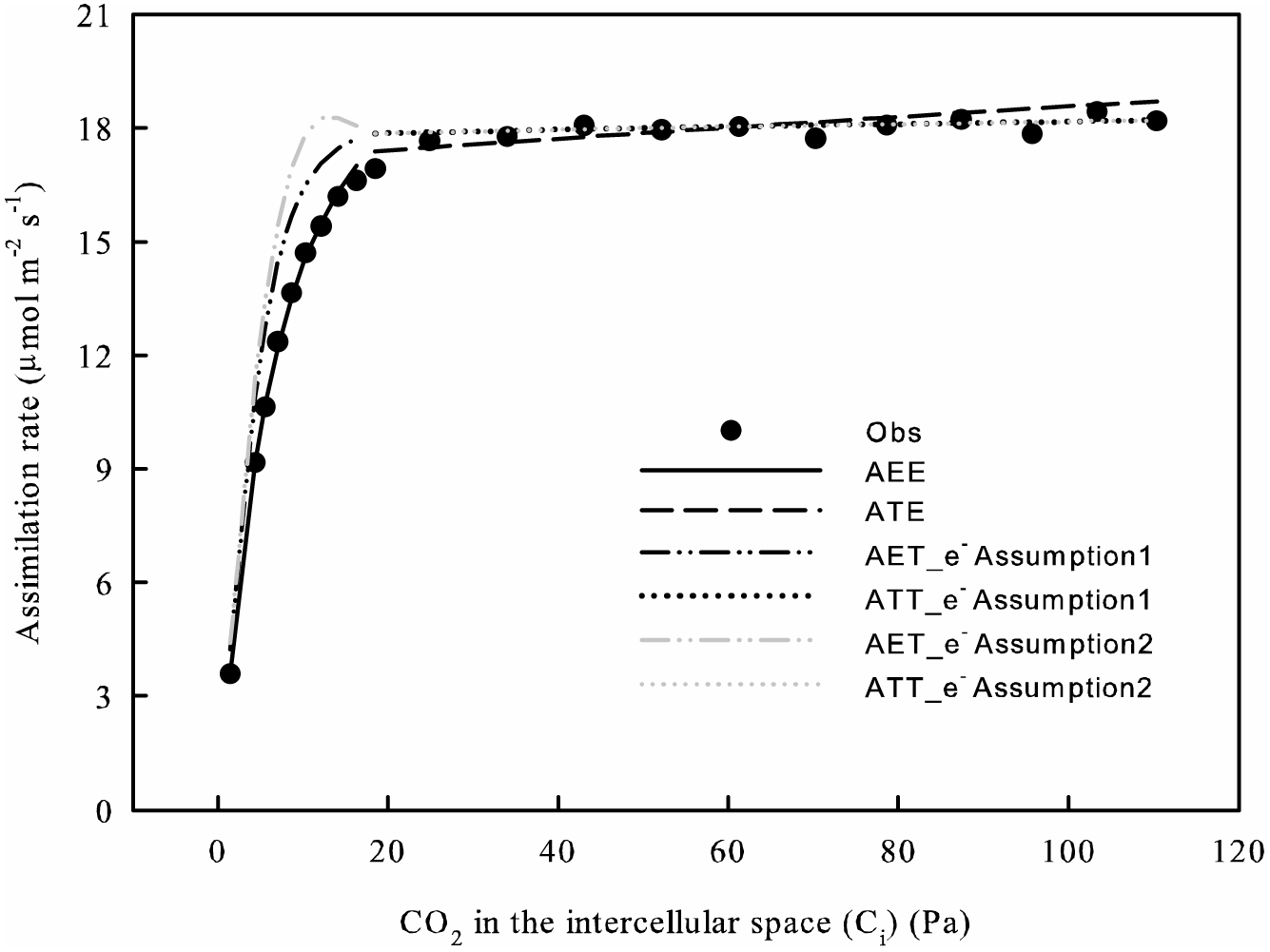
Assimilation-CO_2_ response curves (*A*/*C*_i_) generated using C_4_ photosynthesis of two different assumptions about electron transport. Photosynthetic parameters (*V*_cmax_, *J*, *R*_d_, *V*_pmax_, and *g*_m_) are the same for both assumptions. AET_e^-^Assumption1 and ATT_e^-^Assumption1 represent results of the assumption that no matter how much electron transport is used for PEP carboxylation/regeneration, a certain amount (x*J*) is confined for this use. AET_e^-^Assumption2 and ATT_e^-^Assumption2 represent results of the assumption that electron transport can be freely distributed between PEP carboxylation/regeneration and RuBP regeneration. Parameters are estimated from *A*/*C*_i_ curve of *T. dactyloides* under the light intensity of 1500 μmol m^-2^ s^-1^. AEE and ATE are the same for both assumptions.

### 3.2 Sensitivity analysis

The parameters *K*_c_, *K*_o_, *γ*\*, *K*_p_, *α*, and *g*_bs_ can vary among species in nature (Cousins et al., 2010) and it is therefore important to know how sensitive our results are to variation in these parameters. We conducted a sensitivity analysis for variation in these parameters on the estimated *V*_cmax_, *J*, *R*_d_, *V*_pmax_, *g*_m_ and *k*_CA_ (Fig. 3). This analysis shows all the estimated parameters are robust under the variation of *α* (Fig. 3A) and showed little variation responding to the change of *γ*\* (Fig. 3E) and *K*_o_ (Fig. 3C); however, the estimated parameters are dependent on the other input parameters to different extents (Fig. 3B, D, F). We calculate the average percentage change of estimated parameters along with the 50 % decrease and 100 % increase of the input parameters. *V*_cmax_ showed some medium extent of sensitivity for *K*_c_, *K*_p_, and *g*_bs_ with the average percentage change of 23.11, 7.54 and 17.69 % respectively. *J* is robust in the variations of *K*_c_, and *g*_bs_ (the average change is less than 2%) and with a medium 6.96 % change for *K*_p_. *k*_CA_ is robust in the variations of *K*_c_, *K*_p_, and *g*_bs_ (average change less than 5%). *V*_pmax_ is sensitive for *K*_p_ with the average change of 27.34%, moderately sensitive to the change of *g*_bs_ with 4.01 % and 13.38% change and is robust for *K*_c_. *R*_d_ is sensitive to *K*_c_, *K*_p_, and *g*_bs_ with the change of 6.73, 43.88 and 13.38%. *g*_m_ is strongly sensitive to *K*_c_, *K*_p_, and *g*_bs_ with the average percentage changes of 22.95, 107.04 and 23.19 %. This results suggest that *V*_cmax_, *J*, *V*_pmax_, and *k*_CA_ estimated using our method are relatively robust.

**Fig. 3.**
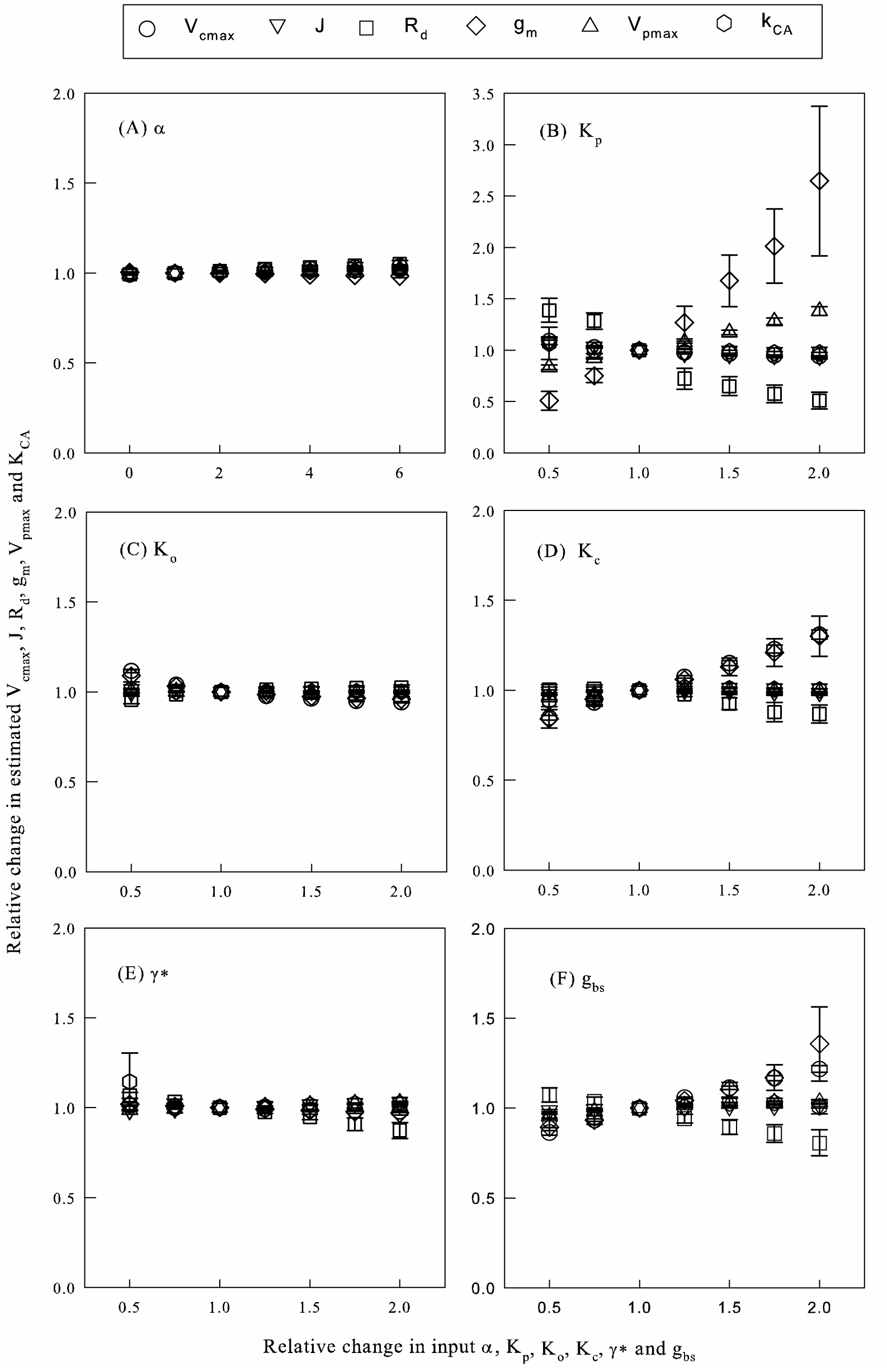
Sensitivity analysis of six estimation parameters to the variation in six input parameters using the model with carbonic anhydrase. Relative changes in the estimated *V*_cmax_, *J*, *R*_d_, *V*_pmax_, *g*_m_ and *k*_CA_ in response to the relative change of six input parameters [(A) *α*, (B) *K*_p_, (C) *K*_o_, (D) *K*_c_, (E) *γ*\* and (F) *g*_bs_] from the initial values in Table S1. The relative change of estimated parameters refers to the ratio of estimated values at a changed input parameter to the estimated value at the initial value of that input parameter. The symbols represent the average change of the nine C_4_ species and error bars represent standard error.

### 3.3 Physiological significance for assimilation rate of the input parameters

In addition to the sensitivity analysis, we performed a simulation analysis to illustrate the physiological importance of input parameters further, and to indicate further the importance of physiological properties in maintaining the efficiency of C_4_ photosynthesis pathway. We chose the estimation parameter set of *T. dactyloides* as an example, held photosynthetic parameters constant *V*_cmax_ (28 μmol m^-2^ s^-1^), *J* (134 μmol m^-2^ s^-1^), *R*_d_ (0.78 μmol m^-2^ s^-1^), *g*_m_ (30.00 μmol m^-2^ s^-1^ Pa^-1^) and *V*_pmax_ (41.91 μmol m^-2^ s^-1^), while changing the values of *α*, *γ*\*, *g*_bs_, and *K*_p_ (as half or twice of the original parameters) to see their effects on the assimilation rate, *C*_bs_ and the O_2_ concentration in bundle sheath (*O*_bs_) (Fig. 4, Table 1). Using photosynthetic parameter sets of other species to perform the simulation analysis yielded similar results (data not shown). The change of α did not lead to changes in assimilation rate (Fig. 4A) and led to small changes in *O*_bs_ (Table 1). The decrease of *γ*\* to half of the current value led to a small change of *C*_bs_ and assimilation rate (less than 0.5 μmol m^-2^ s^-1^) while doubling *γ*\* led to a larger, but still not significant, change (less than 1 μmol m^-2^ s^-1^) (Fig. 4B, Table 1). Importantly, the changes of assimilation rates were less than 0.3 μmol m^-2^ s^-1^ when C_i_ was less than 20 Pa, which is the regular range of *C*_i_ under current ambient CO_2_. However, the change of *g*_bs_ significantly changed the assimilation rate and *C*_bs_ (Fig. 4C, Table 1). The change of *K*_p_ significantly affected the assimilation rate and *C*_bs_ to a large degree under low *C*_i_ (Fig. 4D, Table 1).

**Table 1.** The average change of percentage of CO_2_ concentration (*C*_bs_) and O_2_ concentration at bundle sheath (*O*_bs_) compared to the reference value of *α*0, *γ*\*0, *g*_bs_0 and *K*_p_. Simulation results are obtained by using the original parameter set of *T. dactyloides* with *V*_cmax_ = 28 μmol m^-2^ s^-1^, *J* = 134 μmol m^-2^ s^-1^, *R*_d_ = 0.78 μmol m^-2^ s^-1^, *g*_m_ = 30.00 μmol m^-2^ s^-1^ Pa^-1^ and *V*_pmax_ = 41.91 μmol m^-2^ s^-1^. The values represent average change of percentage of 21 values from 0-120 Pa of intercellular CO_2_ (*C*_i_) (data show mean (standard error)).

| Parameters | $\alpha = 0$ | $\alpha = 2 \alpha 0$ | $\gamma^* = 0.5 \gamma^* 0$ | $\gamma^* = 2 \gamma^* 0$ |
| --- | --- | --- | --- | --- |
| Chang of $C_{bs}$ (%) | -0.91(0.06) | 0.97(0.06) | -2.96(0.28) | 5.05(0.49) |
| Chang of $O_{bs}$ (%) | -6.07(0.30) | 6.01(0.30) | 0.07(0.01) | -0.21(0.02) |
| Parameters | $g_{bs} = 0.5 g_{bs} 0$ | $g_{bs} = 2 g_{bs} 0$ | $K_p = 0.5 K_p 0$ | $K_p = 2 K_p 0$ |
| Chang of $C_{bs}$ (%) | 56.99(3.03) | -29.48(0.41) | 43.12(10.75) | -36.57(4.07) |
| Chang of $O_{bs}$ (%) | 6.77(0.29) | -3.41(0.16) | 0.91(0.18) | -1.18(0.14) |

**Fig. 4.**
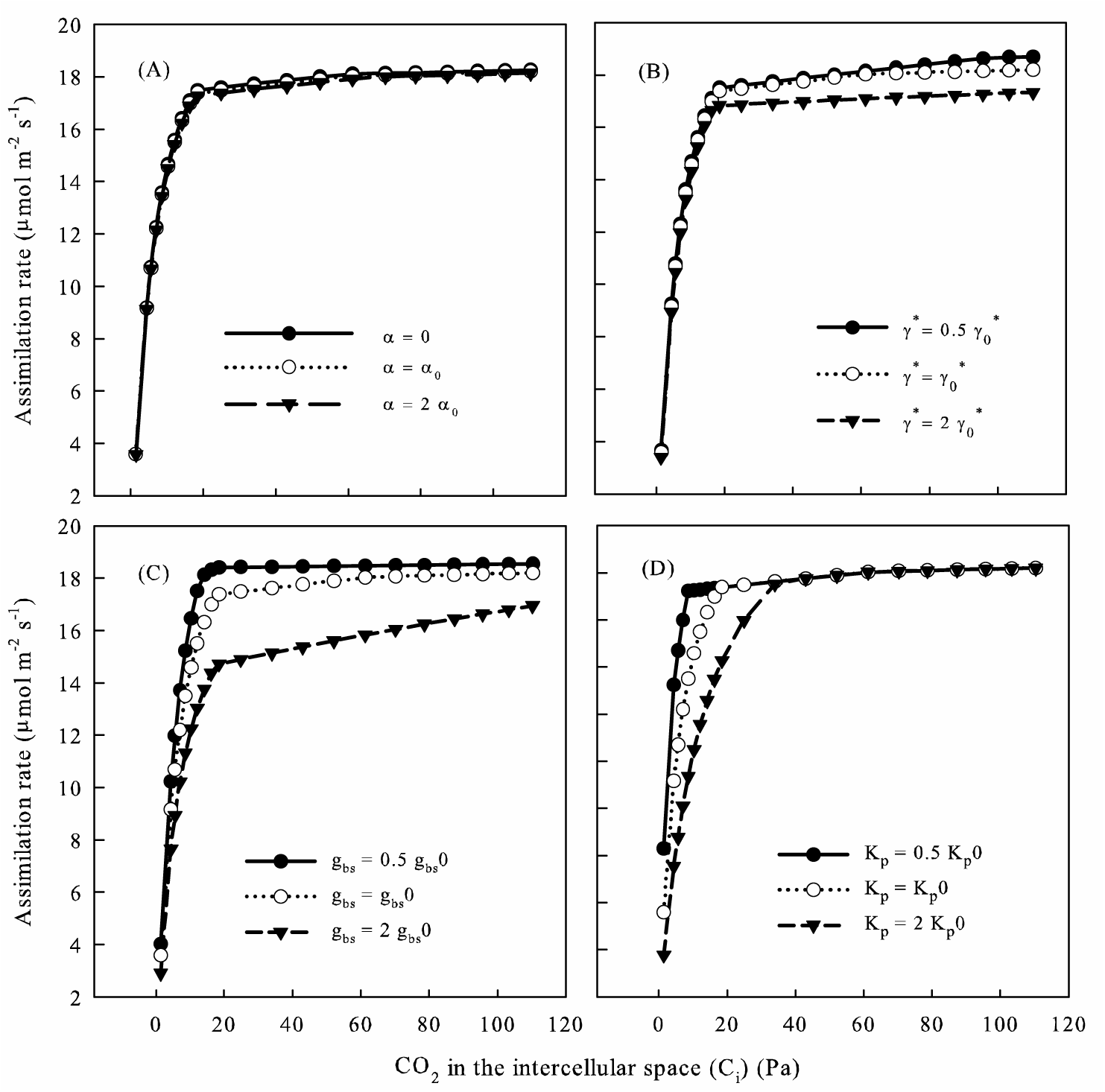
Simulation results of assimilation rate along with different intercellular CO_2_ concentration (C_i_) with the known photosynthetic parameters, but with the change of (A) *α*, (B) *γ*\*, (C) *g*_bs_ and (D) *K*_p_. The original data set are *V*_cmax_ = 28 μmol m^-2^ s^-1^, *J* = 134 μmol m^-2^ s^-1^, *R*_d_ = 0.78 μmol m^-2^ s^-1^, *g*_m_ = 30.00 μmol m^-2^ s^-1^ Pa^-1^ and *V*_pmax_ = 41.91 μmol m^-2^ s^-1^. The reference value of changing parameters at 25°C: *α*0(25) = 0.15, *γ*\*0(25) = 0.000244, *g*_bs_0(25) = 0.0295 and *K*_p_(25) = 8.55 Pa.

### 3.4 Validating the estimation methods

In order to test our estimation methods, we first conducted a simulation test with manipulated error terms. We use the estimated results of the nine species as known parameters (the known values in Fig. 5) to generate new datasets using the C_4_ photosynthesis equations based the first assumption of electron transport and adding error terms to the assimilation rates. The error terms were randomly drawn from a normal distribution of mean zero and standard deviation of 0.1 or 0.2 in an effort to simulate the inevitable random errors in the real measurements. Estimating simulated data sets gave us an idea about how likely we can capture the real parameters of the species given unavoidable errors in measurements. The results show that robust estimation results for *V*_cmax_, *J*, *V*_pmax_, and *R*_d_ can be obtained (Fig. 5A, B, C, D). However, some estimation results of *g*_m_ and *k*_CA_ show some deviation from the real values (Fig. 5E, F).

**Fig. 5.**
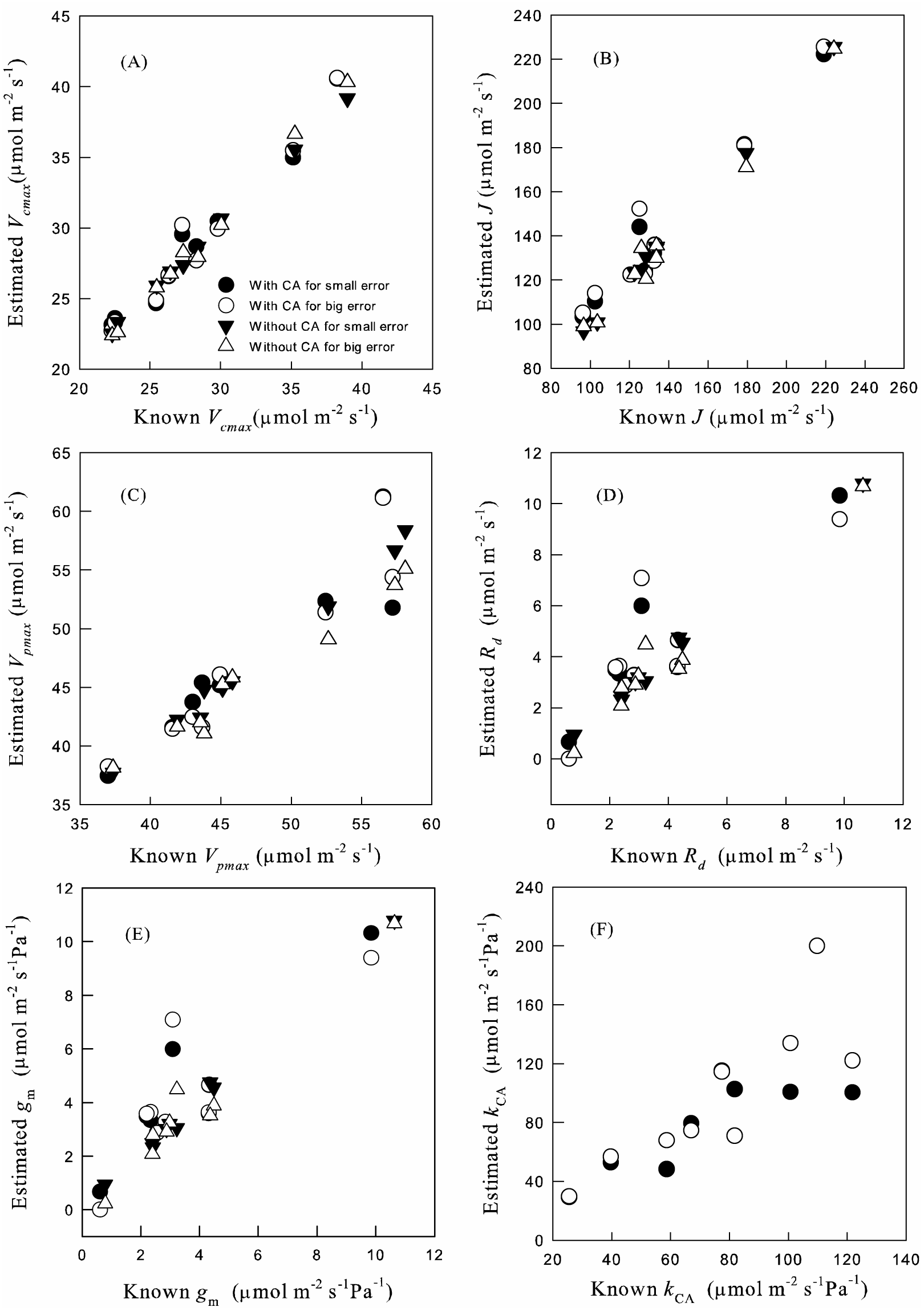
Simulation tests for the estimated parameters ((A) *V*_cmax_, (B) *J*, (C) *V*_pmax_, (D) *R*_d_, (E) *g*_m_ and (F) *k*_CA_) using estimation methods with and without carbonic anhydrase reaction (With CA and Without CA). Datasets are generated by adding random errors for the modeling results using the known photosynthesis parameters of nine species. These known photosynthesis parameters are the true values in the x-axis and are used to compare with the newly estimation parameters. The small error refers to error term randomly chosen with mean 0 and standard deviation of 0.1 and the bigger error refers to error term with randomly chosen mean 0 and standard deviation of 0.2.

To test whether our estimation method could give accurate predictions across typical prediction scenarios, (CO_2_ ranging from 20 Pa to 60 Pa), we performed out of sample tests for our nine target species. To perform these tests, we removed five points of CO_2_ concentrations between 20 and 60 Pa range out of the *A*/*C*_i_ curves and used the rest of the *A*/*C*_i_ curves to estimate parameters. And then we used these parameters to predict the assimilation rate under the CO_2_ concentrations we took out before and calculated the estimation errors. In general, the estimation errors for all our species were small (Table 2).

**Table 2.** Out of sample test results. Five measured points from 20 Pa-60 Pa were taken out when we conducted the estimation process. Then the calculated assimilation rates under these five CO_2_ concentrations were compared with the measured ones. The data shows estimated error between the calculated and measured assimilation rates (data show mean (standard error)).

| Species | *A. virginicus* | *Z. mays* | *E. trichodes* | *P. virgatum* | *P. amarum* |
| --- | --- | --- | --- | --- | --- |
| Model without CA | 0.069(0.036) | 0.150(0.056) | 0.035 (0.017) | 0.193(0.063) | 0.055(0.034) |
| Model with CA | 0.066(0.043) | 0.154(0.057) | 0.111 (0.058) | 0.195 (0.061) | 0.054(0.033) |

| Species | *S. scoparium* | *S. faberi* | *S. nutans* | *T. dactyloides* |
| --- | --- | --- | --- | --- |
| Model without CA | 0.023(0.010) | 0.114(0.055) | 0.258(0.080) | 0.199(0.090) |
| Model with CA | 0.105(0.034) | 0.068(0.040) | 0.263(0.133) | 0.200(0.090) |

We tried to compare our estimation methods with in vitro measurements or other estimation methods using isotopic analysis, especially for *Zea*. Our estimation results for *Zea* obtained similar V_cmax_ with the in vitro estimated Rubisco activity of Pinto et al. (2014); however, the estimated value for V_pmax_ is a little lower than the in vitro PEPC activity measurement with a difference of around 10 μmol m^-2^ s^-1^. For species of the Panicum family with NAD-ME subtype, *P. virgatum* and *P. amarum* in the current study and *P. coloratum* in Pinto et al. (2014), the estimated V_cmax_ and V_pmax_ are quite consistent with the in vitro measurements. Ubierna et al. (2017) reported the g_m_ for *Zea* ranged from 1.69 ± 0.17 to 8.19 ± 0.80 μmol m^-2^ s^-1^ Pa^-1^ using ^18^O and in vitro V_pmax_. Our estimation method fitted a g_m_ for *Zea* of 7.34 μmol m^-2^ s^-1^ Pa^-1^, which falls into the range of their measurements. Barbour et al. (2016) reported a little lower mesophyll conductance for *Zea* using ^18^O measurements.

### 3.5 Validating transition point range

We used chlorophyll fluorescence measurements from seven C_4_ species to test whether the upper and lower boundary CO_2_ concentrations, *CaL* and *CaH*, are reasonable (Table 3). The apparent quantum efficiency of PSII electron transport was calculated with Δ*F/F*m′ = (*F*m′ −*F*s) *F*m′ (Genty, Briantais & Baker 1989). Fluorescence analysis (Baker et al. 2007) is a powerful tool for identifying the limitation states of C_3_ species (Sharkey et al. 2007). If Chlorophyll fluorescence is increasing with increasing CO_2_, *A*_n_ is limited by Rubisco carboxylation limited; when Chlorophyll fluorescence stays constant with increasing CO_2_, *A*_n_ is limited by RuBP regeneration. For C_4_ species, however, the situation is more complicated. Since *V*_p_ could be limited by *V*_pr_ and *V*_pc_ (eq. (9)). Part of the RuBP carboxylation limited condition and RuBP regeneration limited condition for the C_3_ cycle will mix together, leading to a linear increase of fluorescence with increasing of CO_2_, but of a small slope (Fig. S2). Thus, we can only obtain two boundaries of CO_2_ concentrations. Below the lower boundary, *A* and fluorescence increases with increasing *C*_i_ with a steep slope and *A* is RuBP carboxylation and PEP carboxylation limited (AEE); above the higher boundary, *A* and fluorescence is relatively constant along with the increase of *C*_i_ and *A* is RuBP regeneration and PEP regeneration limited (ATT). We measured fluorescence to test whether the upper and lower boundary CO_2_ concentrations, *CaL* and *CaH*, are reasonable. It seems all the *CaL* are above 14 Pa and all the *CaH* are below 65 Pa (Table 3). These results suggest that 10Pa-65Pa is a reasonable range for the transitional point.

**Table 3.** CO_2_ concentration boundaries result for assimilation-limited conditions from fluorescence measurements for seven species. Low: CO_2_ concentration under which assimilation rate increases greatly with increasing CO_2_ (potentially assimilation is limited by PEP carboxylation and RuBP carboxylation). High: CO_2_ concentration above which assimilation rate no longer increases with increasing CO_2_ (potentially assimilation is limited by PEP regeneration and RuBP regeneration). Data show the mean (standard error).

| Species | *P. virgatum* | *P. amarum* | *S. scoparium* | *S. nutans* |
| --- | --- | --- | --- | --- |
| Low(Pa) | 14.1(1.12) | 18.0(1.09) | 17.8(1.09) | 17.6(0.28) |
| High(Pa) | 34.1(1.78) | 55.5(1.40) | 53.1(1.10) | 63.1(2.07) |
| Species | *T. dactyloides* | *T. flavus* | *B. mutica* |  |
| Low(Pa) | 13.8(0.35) | 14.9(2.35) | 15.8(1.13) |  |
| High(Pa) | 46.1(0.20) | 41.4(1.73) | 42.3(1.24) |  |

## 4. DISCUSSION

The photosynthetic parameters from the estimation method are good indicators for the biochemical and biophysical mechanisms underlying the photosynthesis processes of plants. Together with photosynthesis models, they can provide powerful information for evolutionary and ecological questions in both physiological and ecosystem response to natural environmental variation and climate change, to illustrate evolutionary trajectory of C_4_ pathway, as well as in efforts to improve crop productivity (Osborne & Beerling, 2006; Osborne & Sack, 2012; Heckmann et al., 2013). Photosynthetic parameters represent different physiological traits, and comparison of these parameters within a phylogenetic background could help us to understand the further divergence of lineages and species through evolutionary time. Additionally, the response of productivity and carbon cycle of vegetation towards the future climate change depends heavily on photosynthesis parameter estimation as input parameters.

Each of the two different fitting procedures has advantages and disadvantages. Yin’s method (Supplementary material II) uses the explicit calculation of assimilation rate and consequently gives lower estimation error. However, it needs a more accurate assignment of limitation states, especially at the lower end. Thus, Yin’s method will be preferable if one has additional support (e.g. fluorescence measurement) to define the limitation states; otherwise, the Yin’s method may give unbalanced results (Fig. 3). However, Sharkey’s method (Supplementary material I) usually can avoid unbalanced results even without ancillary measurements. Thus, it is better to use both procedures to support each other to find more accurate results. For example, one can first use Sharkey’s method to get estimation results and limitation states, and then use them as initial values for Yin’s method.

Our estimation methods yielded similar results when using models with and without carbonic anhydrase reaction processes. Although carbonic anhydrase activity may well be a limiting step for C_4_ cycle (von Caemmerer et al., 2004; Studer et al., 2014; Boyd et al., 2015; Ubierna et al., 2017), its limitation did not greatly affect assimilation rates in this study. Including the carbonic anhydrase reaction makes the model more complex and difficult to get an explicit solution; therefore, the model without carbonic anhydrase could be used as a simplified form yielding flawed but ‘nearly correct’ predicted values as a part of larger models. However, carbonic anhydrase limitation of C_4_ photosynthesis needs the further assessment from physiological or biochemical perspectives, and our estimation method provides another way to derive carbonic anhydrase parameters, which were comparable with *in vitro* measurements (Boyd et al., 2015). In addition, our results for models with and without carbonic anhydrase activity support the proposition of Cousins et al. (2007) that carbonic anhydrase activity may not be a limiting factor for *A*/*C*_i_ curves of C_4_ plants.

Our results show that despite a clear difference between the assumptions of how the products of electron transport are distributed, the results were similar and comparable with studies using different models under measurements of high light intensity. The bump in the second model happens in AET. In AET, assimilation is limited by RuBP regeneration and PEP carboxylation; therefore, PEP regeneration is not reaching *V*_pr_, and the extra electron transport in PEP regeneration could be freely assigned to RuBP regeneration. This effect will weaken as PEP carboxylation increases. However, under lower photosynthetic photon flux density, assimilation rate will be limited more by electron transport, and the separate assumptions concerning electron transport may start to show divergent results.

The photosynthetic parameters from the estimation method used together with photosynthesis models can provide information and inspiration about the evolutionary and physiological importance of different aspects of the C_4_ syndrome (Osborne & Sack, 2012; Heckmann et al., 2013), which can be investigated by empirical measurements. Several examples emanate from our simulation analysis: (1) *α* represents the fraction of O_2_ evolution from photosynthesis occurring in the bundle sheath cells (eq. (4)) and any *α* > 0 means that O_2_ will accumulate in the bundle sheath cells, due to low *g*_bs_ Both the sensitivity analysis and the simulation analysis showed the change of α did not affect the estimated parameters and assimilation rates, because the high *C*_bs_ created by C_4_ carbon concentrating mechanism overcame any increase of *O*_bs_ and did not lead to high photorespiration. Thus, the compartmentation of O_2_ evolution may not have played an important role in the evolution of C_4_ photosynthesis. (2) A lower Rubisco specificity factor (*γ*\*;eq. (11)) means lower specificity for O_2_, higher specificity for CO_2_, and lower photorespiration. In C_3_ species, selection for Rubisco with lower specificity to O_2_ and high specificity of CO_2_ can increase the carbon gain. However, there is a trade-off between the specificity of Rubisco for CO_2_ and its catalytic rate (Savir et al., 2010; Studer et al., 2014). Based on this trade-off, we can hypothesize that since C_4_ elevates CO_2_ around Rubisco relative to the O_2_ concentration, maintaining low specificity might be optimal, in order to get high catalytic rate of the enzyme to reach higher assimilation rate as shown by the empirical measurements of Sage (2002) and Savir et al. (2010). Our simulation analysis showed the increase of specificity for CO_2_ (decrease of *γ*\*) did not increase the assimilation rate much, which indicates the selection upon Rubisco specificity in C_4_ plants should be relaxed. (3) *g*_bs_ represents CO_2_ leakage from bundle sheath to the mesophyll cell, and changes in *g*_bs_ significantly change the assimilation rate and *C*_bs_. Therefore, avoiding CO_2_ leakage was of great importance for the evolution and efficiency of C_4_ photosynthesis pathway (Brown and Byrd, 1993; Ubierna et al., 2013; Kromdijk et al., 2014).

Although we have shown that parameter estimation can be achieved solely with *A*/*C*_i_ curves, it is easy to combine our methods with ancillary measurements to yield more accurate estimation results by defining the parameters as estimated or known or add additional constraints (Supplementary Material IV). Yin et al. (2011b) proposed a method to obtain R_d_ from the fluorescence-light curve, since the method used for C_3_ species, the Laisk method, is inappropriate (Yin et al., 2011a). Additional measurement of dark respiration could be an approximation for *R*_d_ or could help to provide a constraint for estimating *R*_d_ in our estimating method. Ubierna et al. (2017) discussed the estimation method of *g*_m_ using instantaneous carbon isotope discrimination. With external measurement results, one can change estimated parameters (such as *R*_d_, *g*_m_ and *J*) as input parameters, instead of output parameters, in this curve fitting method (Supplementary material IV). Additional methods, such as in vitro measurements (Boyd et al., 2015; Pedomo et al., 2015) and membrane inlet mass spectrometry (Cousins et al., 2010) of *V*_cmax_, *V*_pmax_, and carbonic anhydrase activity can also provide potential parameter values. Furthermore, if some output parameters are determined in the external measurements, one can also relax the input parameters (such as *g*_bs_) and make them estimated parameters (Supplementary material IV).

## 5. Conclusion

We have developed new, accessible estimation tools for extracting C_4_ photosynthesis parameters from intensive A/C_i_ curves. Our estimation method is based on an established estimation protocol for C_3_ plants and makes several improvements upon C_4_ photosynthesis models. External measurements for specific parameters will increase the reliability of estimation methods and are summarized independently. We developed estimation methods with and without carbonic anhydrase activity. The comparison of these two methods allows for an estimation of carbonic anhydrase activity, and further shows that the method that did not consider carbonic anhydrase activity was a sufficient simplification for C_4_ photosynthesis. We tested two assumptions related to whether the electron transport is freely distributed between RuBP regeneration and PEP regeneration or certain proportions are confined to the two mechanisms. They show similar results under high light, but they may diverge under low light intensities. Simulation test, out of sample test, fluorescence analysis, and sensitivity analysis confirmed that our methods gave robust estimation especially for V_cmax_, J, and V_pmax_.

## Author contributions

HZ, EA and BH conceived the ideas, designed methodology, analyzed the data and led the writing of the manuscript; HZ collected the data; HZ and BH coordinate the study. All the authors contributed to the critical review of the manuscript and approved its final version.

## ACKNOWLEDGMENT

We thank Dr. Jesse Nippert, Kansas State University, for providing the fluorometer chamber.

## Abbreviations

*a*: light absorptance of leaf
*A*_c_: Rubisco carboxylation assimilation rate
*AEE*: RuBP carboxylation and PEPc carboxylation limitation assimilation
*AET*: RuBP regeneration and PEP carboxylation limitation assimilation
*A*_g_: gross CO_2_ assimilation rate per unit leaf area
*A*_j_: RuBP regeneration assimilation rate
*A*_n_: net CO_2_ assimilation rate per unit leaf area
*ATE*: RuBP carboxylation and PEPc regeneration limitation assimilation
*ATT*: RuBP regeneration and PEPc regeneration limitation assimilation
α: the fraction of O_2_ evolution occurring in the bundle sheath
*c*: scaling constant for temperature dependence for parameters
*CaL*: Lower boundary CO_2_ under which assimilation is limited by RuBP carboxylation and PEPc carboxylation
*CaH*: Higher boundary CO_2_ above which assimilation is limited by RuBP regeneration and PEPc regeneration
*C*_bs_: bundle sheath CO2 concentration
*C*_i_: intercellular CO_2_ concentration
*C*_m_: mesophyll CO2 concentration
*ΔH*_a_: energy of activation for temperature dependence for parameters
*ΔH*_d_: energy of deactivation for temperature dependence for parameters
*ΔS*: entropy for temperature dependence for parameters
*ϕ*_PSII_: quantum yield
*γ\**(25): the specificity of Rubisco at 25°C
*g*_bs_: bundle sheath conductance for CO_2_
*g*_bso_: bundle sheath conductance for O_2_
*g*_m_: mesophyll conductance for CO_2_
*I*: light intensity
*J*_max_(25): maximum rate of electron transport at 25°C
*K*_c_(25): Michaelis-Menten constant of Rubisco activity for CO_2_ at 25°C
*K*_o_(25): Michaelis-Menten constants of Rubisco activity for O_2_
*K*_p_(25): Michaelis-Menten constants of PEP carboxylation for CO_2_
*O*_bs_: O_2_ concentration in the bundle sheath cells
Q_10_ for *K*_p_: temperature sensitivity parameter for *K*_p_
*R*: the molar gas constant
*R*_d_: daytime respiration
*R*_dbs_: daytime respiration in bundle sheath cells
*R*_dm_: daytime respiration in mesophyll cells
Rubisco: ribulose-1,5-bisphosphate carboxylase/oxygenase
RuBP: ribulose-1,5-bisphosphate
*T*_k_: leaf absolute temperature
*V*_c_: velocity of Rubisco carboxylation
*V*_cmax_(25): maximal velocity of Rubisco carboxylation at 25°C
*V*_p_: PEP carboxylation
*V*_pc_: PEPc reaction rate
*V*_pmax_(25): maximal PEP carboxylation rate at 25°C
*V*_pr_: PEP regeneration rate
*x*: the maximal ratio of total electron transport could be used for PEP carboxylation.

